# Expression and regulation of SETBP1 in the song system of male zebra finches (*Taeniopygia guttata*) during singing

**DOI:** 10.1101/2024.06.05.597622

**Authors:** Dana Grönberg, Sara Luisa Pinto de Carvalho, Nikola Dernerova, Phillip Norton, Maggie M. K. Wong, Ezequiel Mendoza

**Author notes:** Corresponding author email address: Dr. Ezequiel Mendoza. Contribute equally (Maggie M. K. Wong, Ezequiel Mendoza).

## Abstract

Rare *de novo* heterozygous loss-of-function *SETBP1* variants lead to a neurodevelopmental disorder characterized by speech deficits, indicating a potential involvement of SETBP1 in human speech. However, the expression pattern of SETBP1 in brain regions associated with language remains poorly understood, along with the underlying molecular mechanisms linking it to speech. In this study, we examined SETBP1 expression in the brain of male zebra finches, a well-established model for studying vocal production learning. We demonstrated that zebra finch SETBP1 exhibits a greater number of exons and isoforms compared to its human counterpart. We characterized a SETBP1 antibody and showed that SETBP1 colocalized with FoxP1, FoxP2, and Parvalbumin in key song nuclei. Moreover, SETBP1 expression in neurons in Area X is significantly higher in zebra finches singing alone, than those singing courtship song to a female, or non-singers. Importantly, we found a distinctive neuronal protein expression of SETBP1 and FoxP2 in Area X only in zebra finches singing alone, but not in the other conditions. We demonstrated SETBP1’s regulatory role on FoxP2 promoter activity *in vitro*. Taken together, these findings provide compelling evidence for SETBP1 expression in brain regions to be crucial for vocal learning and its modulation by singing behavior.

## Introduction

SETBP1 (SET Binding Protein 1) was first discovered in 2001 as a protein that binds to the SET protein and is involved in tumorigenesis and leukemogenesis when disrupted by somatic missense variants [1-4]. Interestingly, different types of germline genetic disruption in the *SETBP1* gene cause clinically distinct neurodevelopmental disorders with a broad and variable clinical spectrum. Rare *de novo* heterozygous missense variants clustering in a specific degron region in SETBP1 that is important for its degradation cause Schinzel-Giedion syndrome (SGS, MIM #269150, OMIM 269150), a severe multi-system developmental disorder where most affected individuals do not survive infancy [5, 6]. In contrast, heterozygous *de novo* germline loss-of-function (LoF) variants (truncating and *SETBP1*-specific microdeletions) lead to *SETBP1-*haploinsufficiency disorder (MIM #616078, OMIM 606078) which is a milder neurodevelopmental disorder showing a broad range of symptoms with variable severity [7]. The most commonly observed clinical features were developmental delay, speech impairment, intellectual disability, hypotonia, vision impairment, attention/concentration deficits, and hyperactivity [8]. The speech and language phenotypes of individuals with *SETBP1-*haploinsufficiency were systematically characterized in another partially overlapping cohort. The core characteristics include articulatory, spoken, and written language deficits, with childhood apraxia of speech (CAS) as the most common diagnosis [9]. Interestingly, heterozygous pathogenic LoF variants in *SETBP1* have been independently identified by exome/genome sequencing in different cohorts of individuals with CAS, suggesting SETBP1 involvement in human speech and language [10-12]. Moreover, a recent study systematically characterized the clinical and functional spectrum of missense variants within and outside the degron region. The research found that while variants within the degron region primarily caused SGS, variants outside this region cause milder phenotypes (including prominent speech and language deficits) more similar to haploinsufficiency disorder [13], further indicating a potential role of SETBP1 in speech and language.

Zebra finches are thus far the most well-established animal model for studying vocal learning [14, 15]. There are notable similarities between human speech and zebra finch song learning, ranging from behavioral aspects [15] to genetic factors that have been found to correlate with similar deficits [16, 17]. Zebra finch could therefore serve as a suitable model for studying the function of SETBP1 in vocal production learning. Despite SETBP1 being known since 2001 and its association with clinically distinct syndromes more than a decade ago, there are only a handful of studies examining SETBP1 expression [18, 19] or function in the brain [20-22]. In mice, a few articles have been published examining the effects of SGS SETBP1 variants on various tissues [23, 24], but none have specifically focused on the LOF variants. Cross-species analysis is necessary to determine if the evolution of SETBP1 could be related to language evolution. In this study, we therefore systematically examined the expression pattern of SETBP1 in the brains of male zebra finches, a well-established vocal learning model, and investigated the influence of different singing contexts on its expression.

## Results

### The zebra finch SETBP1 gene

We first cloned the complete *SETBP1* cDNA from male zebra finch brain tissue, and revealed that *SETBP1* consists of six exons and results in four isoforms (**Figure 1**, NCBI accession numbers OR257526-OR257529). In contrast to humans and mice, whose *SETBP1* is located in autosomes, zebra finch *SETBP1* is located on the Zq arm of the sex chromosome Z, which means that males have two copies of *SETBP1* (males have sex chromosomes ZZ) while females have one copy (ZW). In addition, zebra finch *SETBP1* has one more coding exon which results in two more isoforms, compared to humans and mice. The nucleotide sequence of the longest isoform (IsoA) contains 4,845 bp and codes for 1,614 amino acids. The amino acid sequence of SETBP1-IsoA is about 75% similar to the human or mouse sequence. The conservation of the protein domains varies from 50% (second PEST domain) to 100% (first two AT-hook domains) compared to humans (supplementary table 1). Most of the protein domains that are present in human SETBP1 are also present in zebra finch SETBP1 with the exception that zebra finch SETBP1 lacks the last two PPLPPP domains (supplementary table 1).

**Figure 1.**
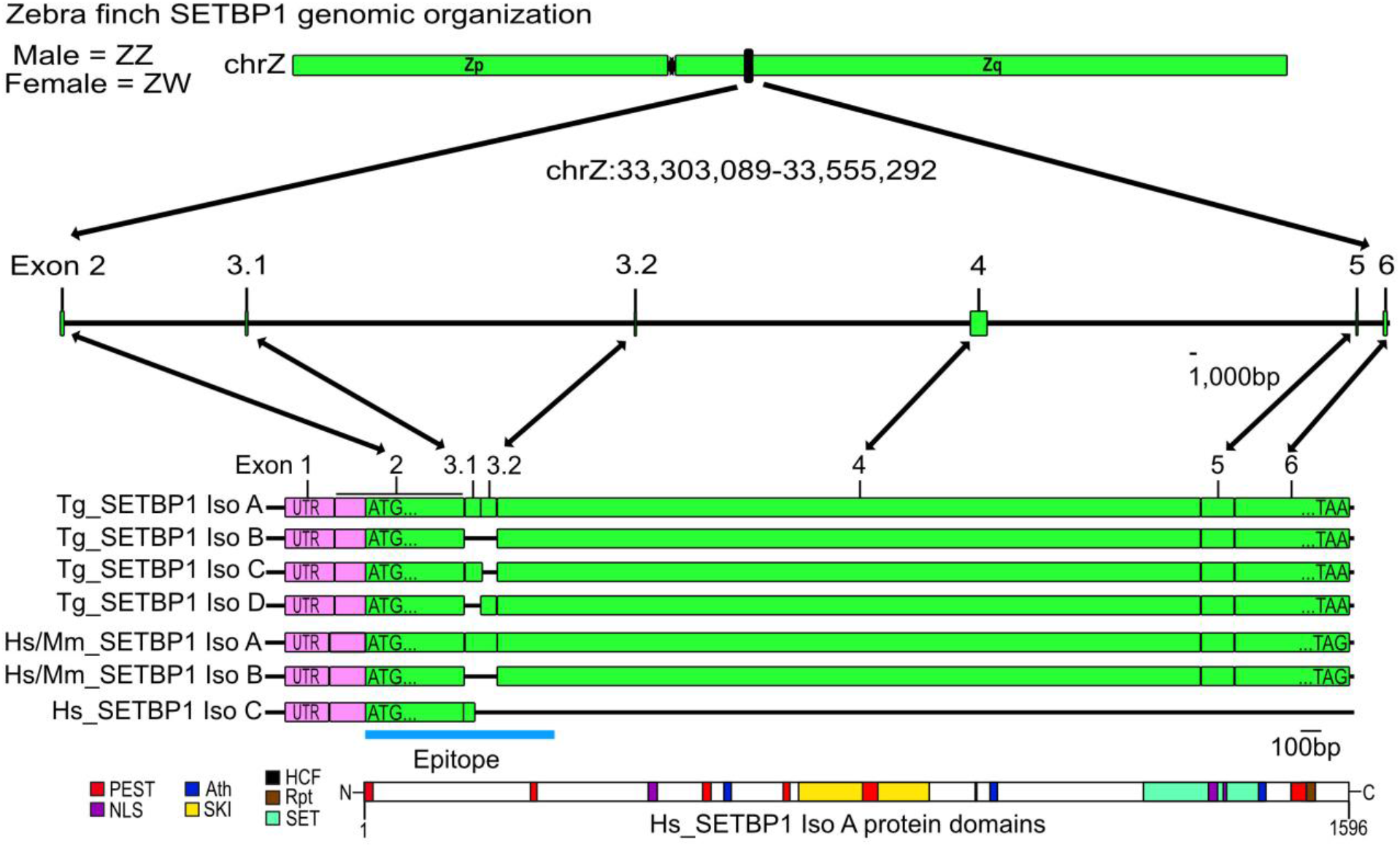
*SETBP1* genomic organization. The SETBP1 zebra finch gene is found in the Z chromosome in the Zq arm. The exact position is shown in black. The first non-coding exon is shown in pink and all coding exons are shown in green. Schematic representations of all isoforms in zebra finches (Tg) and Humans and mice (Hs/Mm) are shown. The sequence of the epitope of the antibody used in this study is depicted with a blue line. Lastly, a schematic representation of the longest human isoform of the SETBP1 protein is shown, with all the known protein domains displayed. Scale bar in the genomic part = 1,000bp and the coding sequence = 100bp.

### A SETBP1 antibody detects both SETBP1 isoforms *in vitro* and in the zebra finch brain

Next, we went on to assess where SETBP1 is expressed in the brain of male zebra finches. Using the protein sequence of zebra finch SETBP1, we identified a SETBP1 antibody with an epitope at the beginning of the protein that was 64.71% similar to the zebra finch SETBP1 protein, 85.19% similar in the sequence of the first coding exon. We then characterized the specificity of this SETBP1 antibody for zebra finch SETBP1, *in vitro* and *in vivo*. We first tested *in vitro* if the SETBP1 antibody would detect the longest and shortest zebra finch SETBP1 isoforms exogenously expressed in HEK cells using immunoblotting (**Figure 2**). Both isoforms of the zebra finch SETBP1 were detected by the SETBP1 antibody, revealing a band of approximately 185kDa, matching the expected size of SETBP1 isoforms (**Figure 2a**). This specific band was not observed in protein lysates of cells expressing only the pMAX vector (GFP transfection control) or the empty vector (EV). Furthermore, immunohistochemical detection in HEK cells expressing the same zebra finch SETBP1 isoforms produced a signal only where SETBP1 was expressed. No signal was observed in cells transfected with an empty vector or in no-primary antibody controls (NPC) (**Figure 2b**). These results suggested that this SETBP1 antibody could detect zebra finch SETBP1 isoforms *in vitro*.

**Figure 2.**
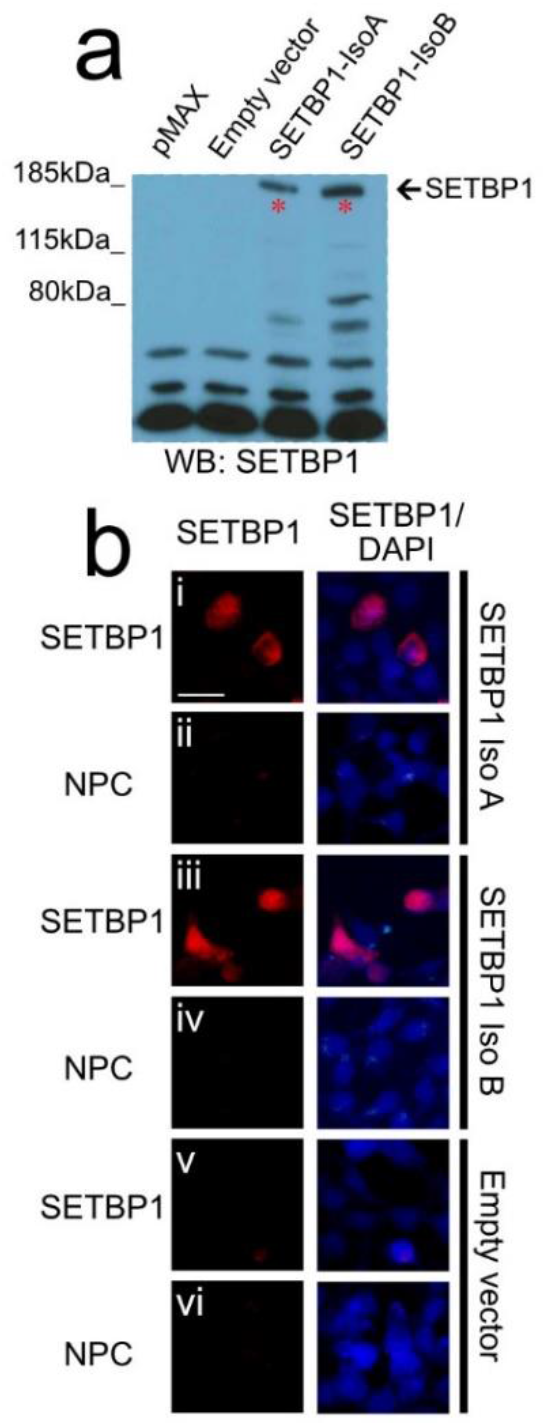
*In vitro* characterization of the SETBP1 antibody PA5-96609 for zebra finches. A) Western blot analysis of protein lysates of HEK cells transfected, from left to right, with pMAX, empty vector, zebra finch SETBP1-isoA and zebra finch SETBP1-isoB and detected with the SETBP1 antibody. A specific band with the expected size was only seen in the lysates that expressed zebra finch SETBP1. B) Immunodetection of SETBP1 in HEK cells transfected with zebra finch SETBP1-isoA (i-ii) and SETBP1-isoB (iii-iv) or with an empty vector (v-vi) and no primary antibody controls (NPC ii, iv and vi). Scale bar= 20µm

To further characterize the specificity of the SETBP1 antibody, we performed immunohistochemistry on brain slices of male zebra finches (**Figure 3b** and **c**). The SETBP1 antibody showed specific detection of SETBP1 in neurons in the brain of zebra finches (**Figure 3a**). In addition, this detection was diminished or abolished when we pre-incubated the antibody with protein lysates of either zebra finch SETBP1-IsoA and -IsoB before proceeding with immunohistochemistry. In contrast, pre-incubation with lysates expressing the empty vector did not affect the detection of SETBP1 (**Figure 3d**). The antibody also detected the shortest isoform B, which lacks exons 3.1 and 3.2 (**Figure 1**). Notably, this region exhibits a high degree of conservation in the amino acid sequence. These findings suggest that the main epitope recognized by the SETBP1 antibody is located in the region of exon 2 (the first coding exon) of zebra finch SETBP1, which is conserved to humans as aforementioned. Taken together, our results demonstrate that the tested SETBP1 antibody detects both isoforms of zebra finch SETBP1 (IsoA and IsoB) using western blot and immunohistochemistry, both *in vitro* and *in vivo*. These findings contribute to the validation of antibody specificity for future studies involving SETBP1 expression and function in zebra finches and potentially other avian species.

**Figure 3.**
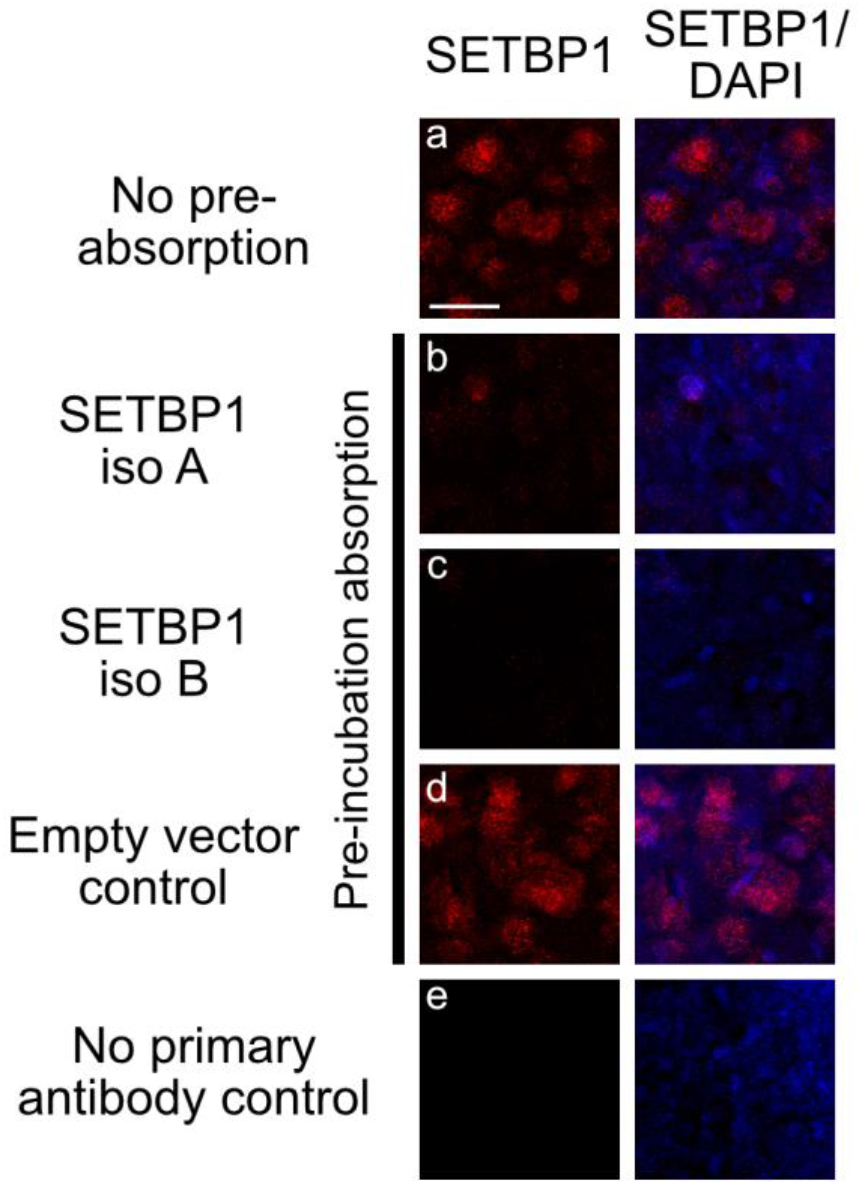
*In vivo* characterization of the SETBP1 antibody for zebra finches. The strong staining (a) was abolished or strongly reduced on brain slices when the SETBP1 antibody was pre-incubated with HEK protein lysates overexpressing SETBP1-Iso A (b) or SETBP1-Iso B (c). This was not the case when lysate of cells transfected with an empty vector was used for pre-incubation (d). No staining was observed in no primary antibody control (e). All photos were taken in mesopallium, a region with a high expression of zebra finch SETBP1 (supplementary figure 1). Scale bar= 20µm

### The SETBP1 protein is prominently expressed in nuclei of the song system in the zebra finch brain

To identify the regions where SETBP1 is expressed, we conducted DAB immunostainings in the brains of non-singing adult males, which serve as a control condition to avoid detecting singing-related expression changes. The DAB staining revealed a homogenous expression of SETBP1 throughout the zebra finch brain (supplementary figure 1). Furthermore, we observed that SETBP1 was prominently expressed in key nuclei of the song system, including HVC (**Figure 4** a and k), the robust nucleus of the arcopallium (RA; **Figure 4** b and l) and Area X (AX; **Figure 4** c and m), contrary to the lateral magnocellular nucleus of the anterior neostriatum (LMAN; **Figure 4** d and n) which showed a weaker expression than the surrounding area. In the majority of the brain regions investigated, SETBP1 exhibited a nuclear expression (**Figure 4** k-n). However, interestingly, we also observed cytoplasmic expression of SETBP1 in cells in the nucleus rotundus (RT), in addition to expression in the nucleus (**Figure 4** o and supplementary figure 2). Intriguingly, in Area X, neurons displayed variable SETBP1 expression, where some neurons showed weak staining while others exhibited strong staining (**Figure 4** m). This pattern could indicate bimodal staining intensities analogous of what had previously been described for the FoxP2 protein in zebra finches, specifically in the medium spiny neurons (MSN) of Area X [25, 26]. Using the SETBP1 antibody, we were able to systematically map the expression of SETBP1 in the brains of male non-singing zebra finches.

**Figure 4.**
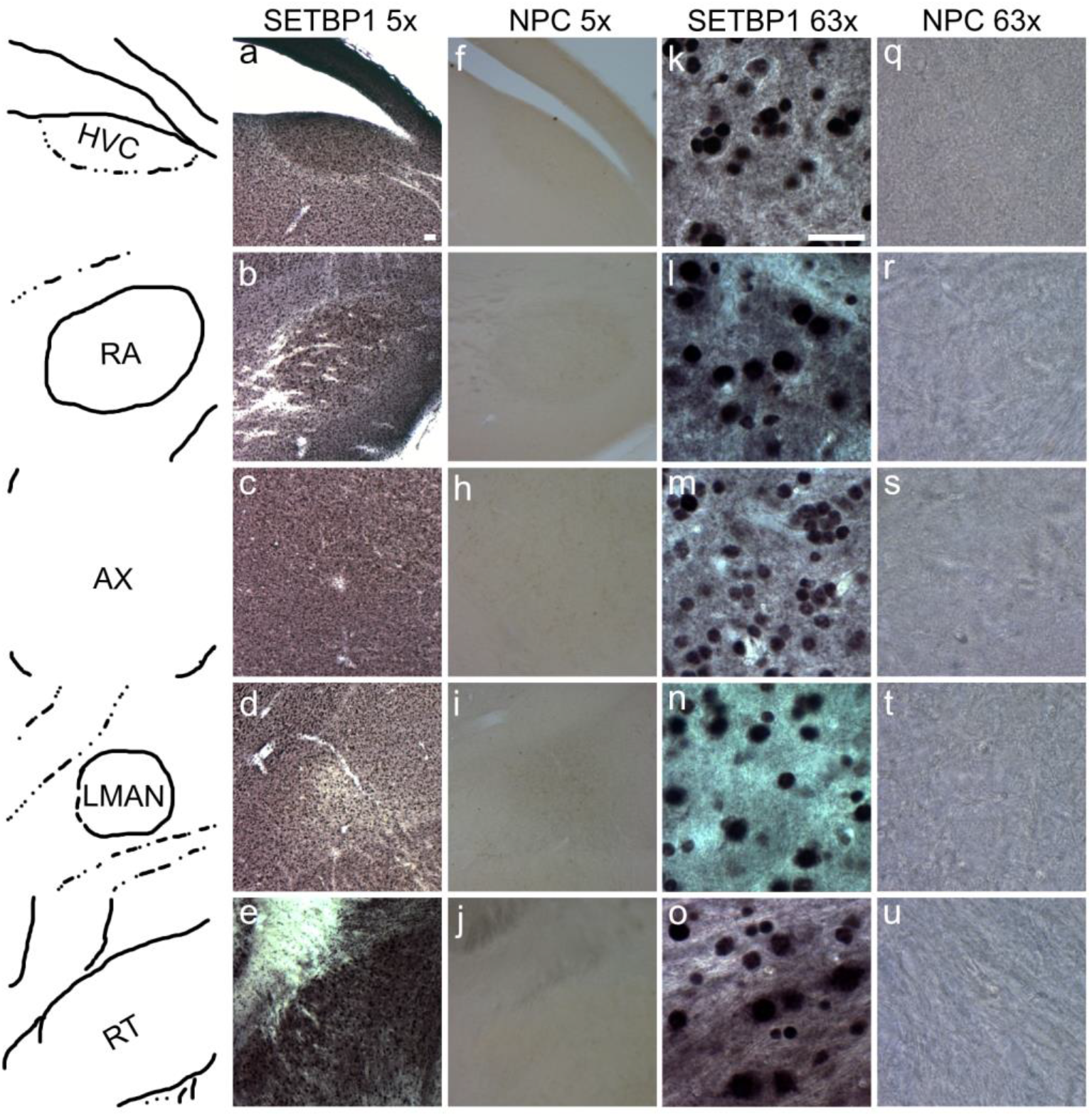
SETBP1 expression in the zebra finch brain. First column shows schematic representations of the different brain regions shown in a-e. DAB immunohistochemistry of SETBP1 on a sagittal slice of a non-singing bird brain showing the regions of HVC (a, f, k and q); the robust nucleus of the arcopallium (RA) (b, g, l and r); Area X (AX) (c, h, m and s); the lateral magnocellular nucleus of the anterior neostriatum (LMAN) (d, I, n and t) and nucleus rotundus (RT) (e, j, o and u) in two different magnifications (5x a-j and 63x k-u) and respective no primary controls (NPC columns f-j and q-u). Scale bar in “a” 5x=50µm and “k” 63x=20µm. Refer to supplementary figure 1 for regions where images (k-u, 63x) were taken.

### The mean intensity of zebra finch SETBP1 in Area X neurons is higher in undirected singers

Previous studies have shown that singing activity and the age of zebra finches can influence the proportion of neurons showing high- or low- FoxP2 intensity in Area X [25]. We therefore investigated whether SETBP1 expression in Area X was affected by these factors. We performed fluorescence immunostainings and confocal imaging, followed by intensity analysis to quantify the intensity levels of SETBP1 (**Figure 5a**). We analyzed the intensities of SETBP1 neurons in male zebra finches under different conditions: undirected singing or singing alone (US, N=7), singing directed to a female (directed singing, DS, N=6), non-singing adults (NS, N=5), and non-singing juveniles (Juvenile, 50 days post-hatching, N=5). Undirected singing is a ‘putative practice state’ of singing while directed singing is a ‘performance state’ in the context of courtship [27]. Interestingly, we did not observe a bimodal pattern of SETBP1 intensities but a normal distribution of intensities ranging from low-to high-SETBP1 expression, in Area X in all of the singing conditions (**Figure 5** b and c), unlike the bimodal pattern described for FoxP2 in zebra finches [25]. The overall mean intensity of SETBP1 in neurons was significantly higher in US birds (94.57±6.322, SEM) compared to NS (63.13±5.537, SEM), DS (62.33±6.085, SEM) and juveniles (58.53±6.987, SEM) (**Figure 5c**; *p<0.05 and **p<0.005, Tukey’s post-hoc test). This suggests that SETBP1 is specifically regulated when birds practice their song (US). Interestingly, we did not find any correlation between singing activity and intensities of SETBP1 in Area X neurons in any of the singing conditions. These results suggest that SETBP1 expression in Area X neurons might vary under different social contexts implying a role of SETBP1 in the neural processes associated with vocal learning and/or production in zebra finches.

**Figure 5.**
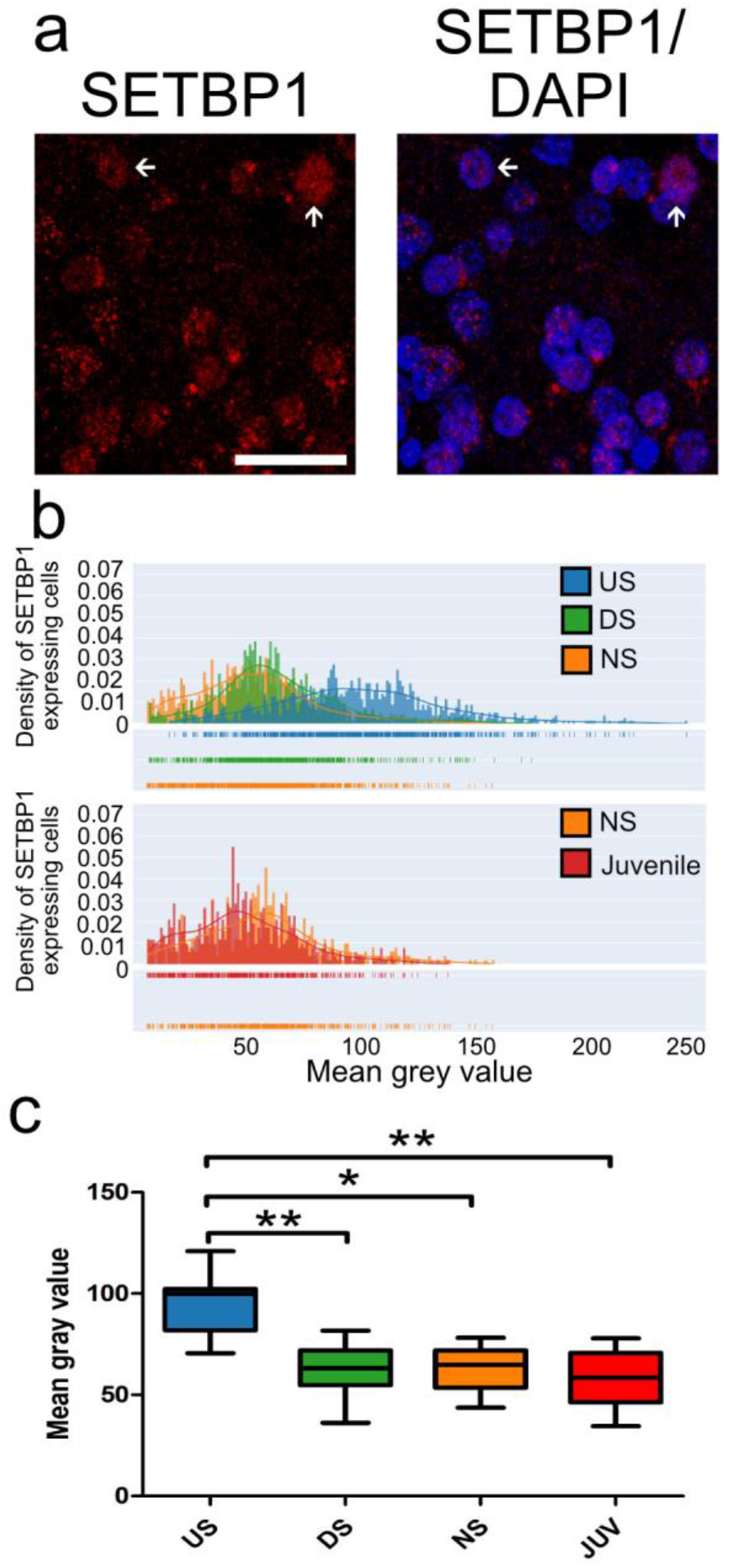
The mean intensity of zebra finch SETBP1 expression in neurons in Area X is higher in undirected singers. a) Example of zebra finch SETBP1 immunostaining in Area X showing weak (arrowhead) and high (arrow) intensities (see also figure 4m). b) Density plots of zebra finch SETBP1 pixel intensities of individual neurons in Area X of undirected singers (blue), directed singers (green), adult non-singers (orange) and juvenile non-singers (50PHDs, red). N = 3200 neurons of 23 zebra finches c) Box plots showing the mean of all zebra finch SETBP1 neuron intensities plotted as a mean for each individual for the different conditions. Sample size: mean of US=7, DS=6, NS=5 and Juveniles=5. *p<0.05; **p<0.005, one-way ANOVA, Tukey’s Multiple Comparison Test. Scale bar: 50µm

### SETBP1 co-localizes with FoxP1, FoxP2, and Parvalbumin in key nuclei of zebra finch song system

To further characterize the SETBP1-expressing neurons in HVC, RA, and Area X, we performed double immunohistochemistry alongside other key genes involved in vocal learning, namely FoxP1 (expressed in HVC, RA and Area X) and FoxP2 (expressed in Area X), which are also recognized markers for medium spiny neurons (MSNs), and parvalbumin (PV; expressed in HVC, RA and Area X), a marker for interneurons. In HVC, we observed that SETBP1 co-localized (defined as the presence of two fluorochromes on the same physical structure in a neuron) with FoxP1 (**Figure 6a**), indicating its expression in projection neurons of HVC [28]. Additionally, SETBP1 also co-localized with PV (**Figure 6b**), indicating that SETBP1 is also expressed in at least PV-positive interneurons in HVC. Similarly, co-localization of SETBP1 with FoxP1 (**Figure 6a**) or PV (**Figure 6b**) was also observed in RA. In Area X, SETBP1 co-localized with FoxP1 (**Figure 6a**), PV (**Figure 6b**), or FoxP2 (**Figure 7a**). We quantified the percentage of neurons showing co-localization of SETBP1 with FoxP1, FoxP2, or PV in four singing conditions: NS, US, DS, and juvenile male zebra finches. The mean co-localization of SETBP1 and FoxP1 in Area X for NS birds was 53.50% ±6.444 (SEM), while it was 36.64% ± 3.245 (SEM) for DS birds, 36.43% ± 5.360 (SEM) for US birds, and 38.51% ± 8.456 (SEM) for juvenile birds (supplementary figure 3a). The mean co-localization of zebra finch SETBP1 and FoxP2 in Area X for NS birds was 21.38% ± 4.506 (SEM), 27.13 % ±2.917 (SEM) for DS birds, 25.92 % ± 2.361 (SEM) for US birds and 20.79 % ± 4.814 (SEM) for juvenile birds (supplementary figure 3b). The mean co-localization of SETBP1 and PV in Area X for NS birds was 7.537% ± 1.324 (SEM), 5.39 % ±1.236 (SEM) for DS birds, 5.626 % ± 1.687 (SEM) for US birds, and 10.85 % ± 4.043 (SEM) for juvenile birds (supplementary figure 3c). The different singing conditions showed similar extent of SETBP1 co-localization with FoxP1, FoxP2 or PV in Area X (supplementary figure 3). These results suggest that SETBP1 is expressed in interneurons and MSNs or projecting neurons in the brain regions we examined, and that the extent of co-localization with FoxP1, FoxP2, or PV remains stable during singing and zebra finch brain development.

**Figure 6.**
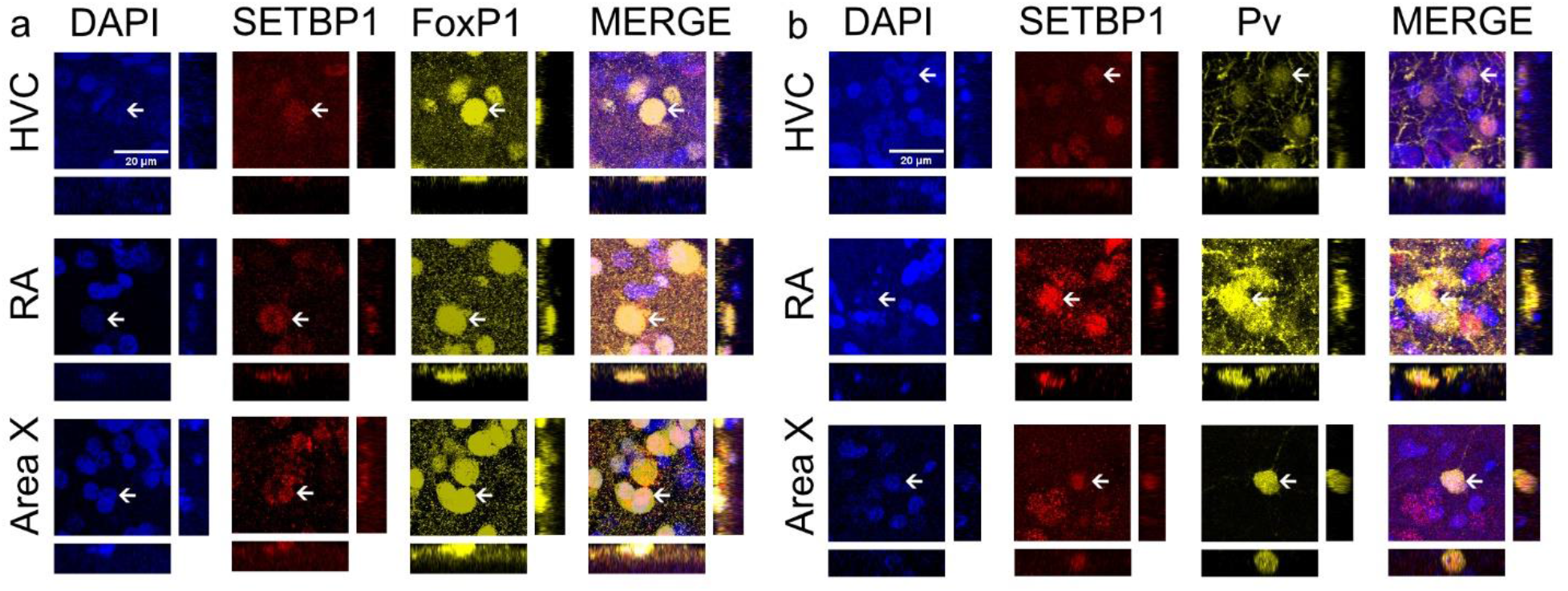
SETBP1 co-localizes with FoxP1 and PV in Area X, HVC and RA. a) SETBP1 (color-coded in red) and FoxP1 (pseudo color-coded in yellow) immunostainings showed colocalization in HVC, RA and Area X. SETBP1 is expressed in projection neurons in HVC and medium spiny neurons in Area X. b) SETBP1 (color-coded in red) and Parvalbumin (pseudo color-coded in yellow) immunostainings showing colocalization in HVC, RA and Area X. SETBP1 is also expressed in interneurons in HVC and Area X. In all areas we show DAPI (color-coded in blue) and merge of all channels. Orthogonal views of the co-localizing cells are show in each panel. Arrows indicate examples of a neuron co-expressing SETBP1 with either FoxP1 or PV. Scale bar = 20µm.

**Figure 7.**
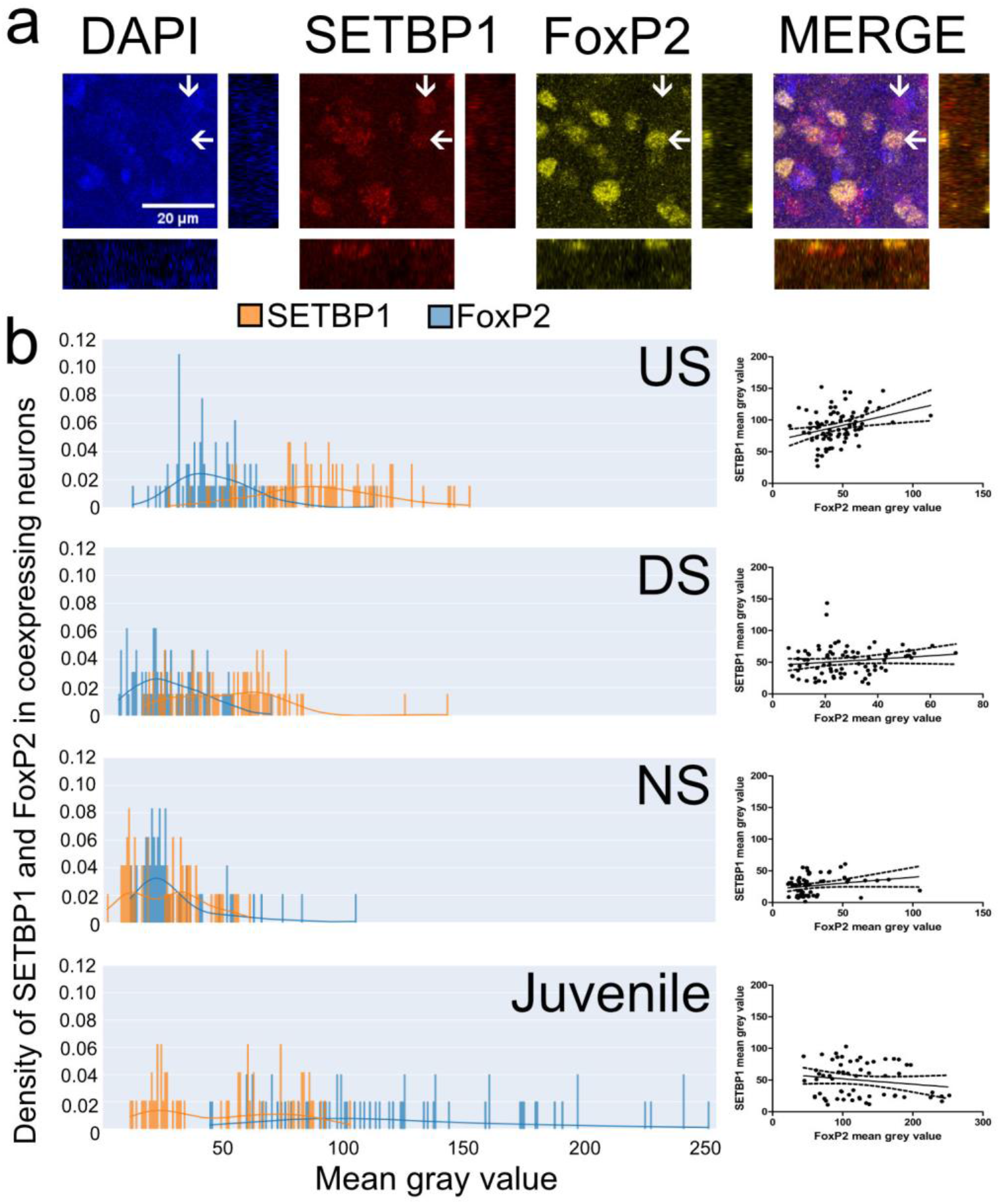
SETBP1 and FoxP2 co-expressing neurons’ intensities correlate with undirected singing. a) DAPI (color coded in blue), SETBP1 (color coded in red) and FoxP2 (pseudo color coded in yellow) immunostaining in Area X and merge of all channels, arrow pointing to the left highlights to a SETBP1+/FoxP2+ example, arrow pointing down a SETBP1+/FoxP2-neuron. Orthogonal views of the co-localizing cell are shown for each channel. b) Density plots of pixel intensities of individual SETBP1 (orange) and FoxP2 (blue) co-localizing neurons in Area X of undirected singers (US), direct singers (DS), non-singers (NS) and 50 PHDs juvenile non-singers (Juvenile). On the right of the density plots we show correlation plots of intensities of SETBP1 and FoxP2 co-localizing neurons in Area X, undirected singers have a significant correlation (Spearman correlation, spearman r= 0.3149, p= 0.0044), all others did not (DS, spearman r= 0.1833, p= 0.1035; NS, spearman r= 0.2264, p= 0.0820; Juveniles, spearman r=-0.09064, p= 0.4910). Scale bar = 20µm

### Expression of SETBP1 and FoxP2 co-localizing medium spiny neurons of Area X changes during development and is induced by undirected singing

We next examined whether there is a correlation between SETBP1 and FoxP2 expression intensities in MSNs expressing both proteins in Area X under different singing conditions (**Figure 7**). Our analyses showed that the fluorescence intensity distributions of SETBP1 and FoxP2 in Area X MSNs displayed different patterns in all singing conditions (**Figure 7b**). US birds showed overall higher mean gray values (MGV), i.e. higher SETBP1 fluorescence intensities while having low-FoxP2 intensities (**Figure 7b**). We only found a statistically significant positive correlation between the two proteins in MSNs in US birds (Spearman correlation, spearman r=0.3149, p=0.0044) but not in the other singing types. In contrast to US, juvenile birds showed an opposite pattern with lower SETBP1 and higher FoxP2 intensities (**Figure 7b**). In DS and NS birds the majority of neurons in which SETBP1 and FoxP2 co-localized showed weak expression of both proteins. Altogether this suggests that SETBP1 and FoxP2 expression correlate in MSNs during US, a period of vocal learning.

### Zebra finch SETBP1 regulates a zebra finch FoxP2 promoter *in vitro*

SETBP1 has been shown to directly regulate two promoters of FOXP2 *in vitro* in human cells [13]. We thus searched for these two promoter regions of FoxP2 in the zebra finch genome. We found two putative regions at similar distances from exon 1 of FoxP2 as described for humans (**Figure 8a**) [29]. The promoter region we identified upstream of TSS1 is 2181 bp long and is 53.16% similar to the human promoter sequence. This promoter sequence is located about 329kb upstream of the coding exon 1 of FoxP2. We identified the zebra finch FoxP2 promoter upstream of TSS2 at about 11kb upstream of the first FoxP2 exon. The promoter region we identified upstream TSS2 is 2501bp long and 81.3 % similar to the human sequence. In addition, two zebra finch 5’UTRs are known in the region of TSS2 (ENSTGUG00000005315:ENSTGUT00000043419.1 and ENSTGUG00000005315:ENSTGUT00000040986.1). We used the promoter regions upstream of TSS1 and TSS2 of zebra finch FoxP2 to drive the expression of Luciferase protein (*Photinus pyralis* synthetic protein) and tested the regulatory effects of all zebra finch SETBP1 isoforms in HEK cells using a luciferase reporter assay. Human SETBP1 was shown to upregulate Luciferase expression via TSS1 and TSS2 of FOXP2 *in vitro* [13]. In contrast to human SETBP1, zebra finch SETBP1 could not regulate FoxP2 TSS1 (**Figure 8b**). Luciferase expression via FoxP2 TSS2 was upregulated by all zebra finch SETBP1 isoforms significantly compared to the empty vector control (one-way ANOVA, followed by Tukey’s multiple comparison test, ∗∗∗p < 0.001; F=50.76). Among zebra finch SETBP1 isoforms, we found that SETBP1-IsoC was the strongest activator of TSS2 (mean of 10.55 ±0.3632, SEM) while SETBP1-IsoA was the weakest activator (7.924 ±0.9074, SEM) (One-way ANOVA, Tukey’s Multiple Comparison Test, ∗p<0.05, q=5.549). Both SETBP1-IsoB (8.732 ±0.1197, SEM) and -IsoD (9.281±0.1552, SEM) led to moderate activation of FoxP2 TSS2.

**Figure 8.**
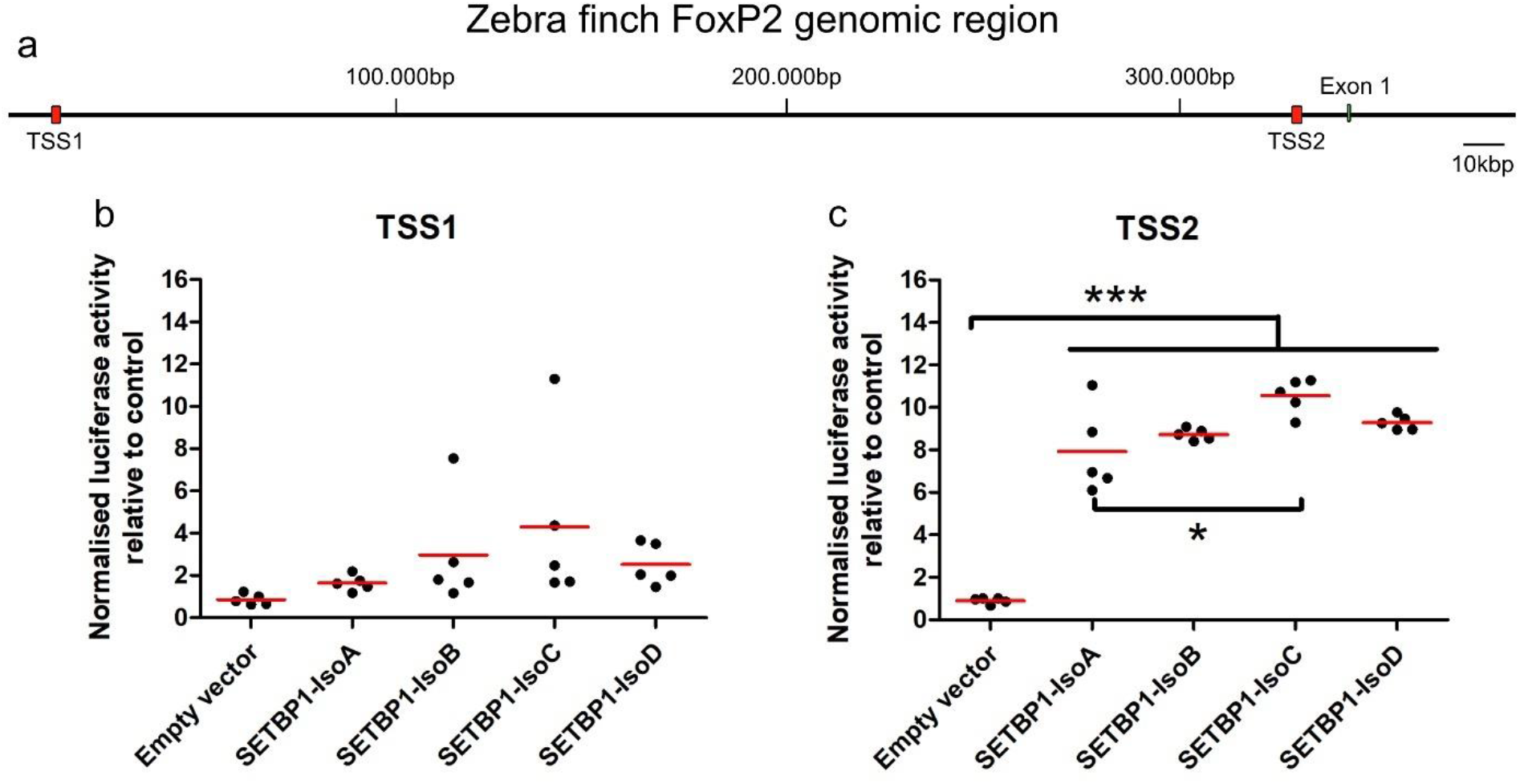
Zebra finch SETBP1 regulates at least one promoter of zebra finch FoxP2 *in vitro*. a) Genomic region of FoxP2 showing two transcription start sites (TSS in red) before the first coding exon of FoxP2 (green) that were used in luciferase assays with overexpression of all zebra finch SETBP1 isoforms. b-c) Dot plot, each dot represents the mean of the luminescence measured from each experiment performed in triplicate, the red line is the mean of means, presented as normalized luciferase activity relative to empty vector control, corrected for transfection by pGL4.75 Renilla luciferase activity. b) Luciferase assays with TSS1 and all SETBP1 isoforms, all were non-significant. c) Significance levels from all SETBP1 isoforms to the empty vector control are represented by stars, ∗0.01 < p <0.05; ∗∗∗p < 0.001. One-way ANOVA; F = 50.76; R squared= 0.9442, DF = 19; n = 5; followed by Tukey’s multiple comparison test; Luciferase assays with TSS2 and all zebra finch SETBP1 isoforms, all activated FoxP2 expression with isoform-C being the strongest activator.

Altogether, our data show that SETBP1 is expressed in regions important for vocal learning in the zebra finch brain, co-localized with proteins whose rare genetic disruptions are correlated with speech and song/vocal learning deficits, and that it can directly regulate FoxP2 expression.

## Discussion

In the present study we systematically examined the SETBP1 expression in regions that are important for vocal learning in the male zebra finch brain. We showed that zebra finch SETBP1 has four isoforms, unlike in humans and mice. We characterized a commercial antibody for its use to detect the SETBP1 protein in the zebra finch brain. We showed that SETBP1 subcellular localization can be either nuclear or cytoplasmic. Furthermore, SETBP1 is expressed in important vocal nuclei in the brain of zebra finches and co-localizes with FoxP1 and FoxP2, whose rare genetic variants are correlated with human speech deficits. Undirected singing activity had the strongest effect on the expression of the SETBP1 protein in Area X, and this was positively correlated with FoxP2 expression and singing dynamics. Finally, we demonstrated the direct regulation of the FoxP2 transcript from TSS2 by all zebra finch SETBP1-isoforms *in vitro*. Overall, our data suggest that SETBP1 and FoxP2 co-localize in MSNs in a key vocal nucleus Area X and that SETBP1 might regulate FoxP2 expression in zebra finches.

Similar to human SETBP1, the zebra finch SETBP1 protein has three nuclear localizing sequences and is predicted to show nuclear expression. However, it was shown that it can also show cytoplasmic expression [1]. In the brain of male zebra finches SETBP1 protein is mostly expressed in the nucleus, but cytoplasmic expression of SETBP1 was also consistently detected in nucleus rotundus. This could suggest a different functional role or regulation of SETBP1 as compared to all other brain regions analyzed where SETBP1 is only found in the nucleus. Human SETBP1 has been shown to function as a transcription factor and directly bind to DNA through the AT hooks and interaction with other proteins [24]. Thus far, only a few SETBP1 interactors have been identified and it remains unclear whether the interaction exists in the brain, let alone regions important for speech development and vocal learning. SETBP1 was first identified as it bound to the SET protein, an oncogene well-studied in non-neural tissues [30]. SETBP1 variants and functional dosage of SETBP1 protein may affect functions of SET such as a) acetylation state of histones which are targets of the INHAT complex; b) phosphorylation state of targets of PP2A protein; c) DNA nuclease activity that may be important in DNA repair; and d) cell cycle through CDKN1A and p53 [31]. All these pathways may affect brain development. SET variants were linked to moderate intellectual disability and in most cases speech delays were reported [32, 33]. Investigating the expression of SET protein in the brain would help to know where SETBP1 and SET may interact or not. Another known binding partner of SETBP1 is HCF1 [34]. *HCF1* variants are linked to intellectual disability and in some cases speech deficiency [35]. However, HCF1 is highly expressed in embryonic brain tissue but not in the adult brain [36], which might suggest that SETBP1 and HFC1 complex could be important during embryonic development but less during a learning context. HFC1 was shown to be expressed in interneurons [37] but we also showed that SETBP1 was not widely expressed in interneurons in zebra finches. This might indicate that interaction with HCF1 might be less relevant for the speech phenotype of SETBP1. Future studies mapping SETBP1 interaction partners will be helpful to understand how SETBP1 and its interactors contribute to vocal learning and speech development.

To date, there are limited studies that analyzed the expression of SETBP1 in the brain. There are thus far two studies showing *SETBP1* expression in mouse cortical cells using single nucleus RNA-seq [18, 23] but this approach does not inform us of spatial localization or abundance of the SETBP1 protein. An individual with SGS was studied with serial MRI showing progressive atrophy in the white matter and especially in the basal ganglia [5, 38]. Other regions shown to be affected were thalamus, brainstem and cerebellum. However, the expression of *SETBP1* in these brain regions have not been investigated. Notably, we found high expression of SETBP1 in all these equivalent regions in the brain of male zebra finches (**Figure 4** and supplementary figure 1). Especially in basal ganglia, we demonstrated SETBP1 expression in MSNs and interneurons in Area X. Of note, SETBP1 expression is higher in undirected singers in MSNs of Area X compared to other singing conditions, implicating its potential role in vocal learning. In Area X, both low- and high-FoxP2-expressing MSNs have been found [25]. FoxP2 expression levels are crucial for vocal learning shown by the fact that downregulation [17, 39] or overexpression [40] during song learning leads to song deficits. Both conditions inhibit dynamic behavioral regulation. Interestingly, we demonstrated that while the number of low-FoxP2-expressing neurons decreased upon singing, the same neurons also showed high SETBP1 level (**Figure 7b**). Using an *in vitro* assay, we showed that SETBP1 can regulate a FoxP2 promoter and its expression. Together, this tight regulation of SETBP1 expression in Area X suggests that the expression levels of SETBP1 might be crucial for song learning in male zebra finches. Further investigation and functional studies will be informative to elucidate the specific role of zebra finch SETBP1 and the genes it regulates and interacts with in the regulation of vocal learning and/or song maintenance in zebra finches.

Overall, this is, to our knowledge, the first systematic characterization of SETBP1 expression in the brains of male zebra finches, a well-established vocal learning model. We showed that SETBP1 is dynamically expressed in song nucleus Area X in male zebra finches and significantly positively correlated with expression of FoxP2, an important regulator of song learning that can itself be regulated by SETBP1. This suggests a potential role for zebra finch SETBP1 in the regulation of vocal learning and/or song maintenance. Our study provides important insights into the zebra finch SETBP1 gene, its expression, and its singing-induced and developmental regulation.

## Materials and Methods

### Animals and brain sectioning

All male zebra finches used in this study were obtained from a breeding colony at the Free Universität, Berlin. The animal husbandry, breeding, and experimental procedures were conducted in strict compliance with the regulations and permits granted by the local Berlin authorities governing research involving animals (TierSchG). For this study, a total of five juvenile zebra finches, aged 50±2 post-hatching days (PHD), and five adult male birds (>100 PHDs) were used. Both of these conditions were non-singing (NS) birds, for which they were recorded for two hours after lights-on, and individuals that sang less than 10 motifs in the last 30 minutes were selected. In addition, we had adult male zebra finches that were chosen if they would sing more than 100 motifs in the last 30 minutes. This was the undirected singer (US) group. Lastly, we had adult zebra finches that were presented with adult females to encourage directed-singing (DS) or courtship song. To ensure that male zebra finches were singing directly to the females, we video-recorded these experiments and exchanged the females every 5 minutes. Again, only birds that sang more than 100 motifs in the last 30 minutes were chosen. Birds that passed these criteria were quickly euthanized by an overdose of isoflurane and transcardially perfused with PBS followed by 4% paraformaldehyde (PFA) in phosphate buffer. Brains were dissected carefully and were kept in 4% PFA overnight for postfixation, then transferred to PBS until they were cut into 70µm thick sagittal sections in the Vibratome (LEICA VT1000S) and stored in wells with PBS until further processing for immunohistochemistry.

### Cloning of SETBP1 cDNAs from zebra finch brain

cDNA from the brains of zebra finches, prepared in our laboratory [41], was used to clone SETBP1. The coding sequence (CDS) of SETBP1 was downloaded from Ensembl (https://www.ensembl.org/index.html) after searching the zebra finch genome for SETBP1 (ENSTGUG00000001615.2). Primers were designed to amplify the entire coding region of zebra finch SETBP1, spanning 4845 base pairs (bp) (first set of primers: 5’-gatGGTACCATGGAGCCCAGAGAGACTTTGAG-3’ forward and 5’-atcGAATTCTTAGGGAAGGCCTTCACTTTCGC-3’ reverse). The forward primer has a KpnI and the reverse an EcoRI restriction site. The resulting polymerase chain reaction (PCR) product using Phusion High-Fidelity DNA Polymerase (Thermo Scientific F-5345) was examined on an agarose gel, cleaned from nucleotides with the Qiaquick PCR purification kit (Qiagen, Chatsworth, CA), and cloned into the pcDNA3.1 vector using the mentioned restriction sites. Initially, we obtained clones with SETBP1-IsoA and -IsoB. To specifically discern between the other isoforms, a series of PCRs with a different set of primers (second set of primers: 5’-ACCACCAAGAGAGCGAAGAA-3’ forward and 5’-CACAGGGAACCCACACTC-3’ reverse; 5’-CCTTGGTGGCACTAATTGCT-3’ forward and 5’-GTGGTTGCAGAAAAGGGAAA-3’ reverse, in both cases resulting in two bands indicating the presence of isoC and isoD in tissue) was done to see if they would express SETBP1 from cDNA. A first amplification round was done with the first set of primers, and clones were picked and genotyped using the second set of primers to select for the remaining isoforms. At least four clones were sequenced for each isoform. The sequences of the four SETBP1 isoforms were deposited to NCBI (NCBI accession numbers: OR257526-OR257529).

### Antibody characterization

The protein sequence of SETBP1 from zebra finches was utilized to search for antibodies that would be compatible. The antibody with the most conserved sequence found in our epitope comparisons was the rabbit polyclonal anti-SETBP1 (Invitrogen, rabbit polyclonal, PA5-96609 batch XC358766A, concentration 1.82 mg/ml, RRID:AB_2808411) with an epitope between amino acids 1-242 of human SETBP1. To characterize this SETBP1 antibody, the zebra finch SETBP1-IsoA (longest) and -isoB (shortest) proteins were expressed in HEK293 cells followed by western blotting, showing a band corresponding to approximately 185 kDa molecular weight in each case. Furthermore, to characterize the antibodies for immunohistochemistry, the two SETBP1 isoforms were overexpressed again in HEK cells and detected with the SETBP1 antibody. Negative controls were prepared by omitting the primary antibodies (NPC) or using empty vectors. In both SETBP1-overexpressing conditions, but not with NPC or empty vector, a signal was detected by immunohistochemistry. Additionally, to block the antibody before incubation on the slides, we preincubated 1µl of the SETBP1 antibody with 25µl of zebra finch SETBP1-isoA or -isoB overexpressing protein lysate in PBS/0.3% Triton-X100 in an ending volume of 500µl, using NPC or 25µl of empty vector lysate as a control. The antibody alone or the antibody with protein lysates were incubated for 120 minutes at 4°C before proceeding with the immunohistochemistry protocol. Only in slices that had no pre-incubation with SETBP1 protein lysate or empty vector lysate did we find a signal, but detection was abolished or diminished in the slices that were pre-incubated with SETBP1 overexpression lysates. The specificity of the FoxP1 antibody (Abcam, mouse monoclonal, ab32010, RRID:AB_1141518) had previously been determined using transient overexpression of human or zebra finch FoxP1 in HEK293 cells and peptide blocking [28, 42]. The specificity of the FoxP2 antibody (Abcam, goat polyclonal, ab1307, RRID:AB_1268914) was characterized for zebra finches by western blot and peptide blocking [25]. The specificity of the parvalbumin (PV) antibody (Swant, mouse monoclonal, PV 235, RRID:AB_10000343) was characterized for zebra finches [43].

### Western blotting

Western blot was conducted following the protocol described previously [42], with the following modifications. Protein concentration was quantified using BCA1 from Sigma. Thirty micrograms of protein lysate were separated by 6% Bis-Tris Gel, then transferred to a polyvinylidene fluoride membrane (Roche, Indianapolis, IN, USA), and blocked with Roti-Immunoblock for 2 hours. The membranes were then incubated with the SETBP1 antibody (dilution 1/2000) overnight at 4°C. Subsequently, the membranes were washed three times with PBS/0.1% Tween 20, followed by incubation with a donkey anti-rabbit IgG POD F(ab’)2 (dilution 1/200,000, Amersham NA9340, RRID:AB_772191) for another 30 minutes. Binding was detected on X-ray films using a Western Lightning Plus Chemiluminescent Substrate detection system for HRP (Perkin-Elmer, Boston, MA, USA, NEL103E001EA).

### DAB-Immunohistochemistry

DAB-immunostaining was performed following the previously described protocol [25], with the following modifications. After washing the slices with PBS containing 3% Triton for 15 minutes, repeated six times, the slices were blocked for 1 hour in ROTI®Immuno Block. Subsequently, the sections were incubated overnight with a primary antibody against SETBP1 (dilution 1:500) in 0.1% Triton/0.1M PBS, applied to the slices at 4°C overnight. For the secondary antibody, a goat anti-rabbit (Vector Laboratories, Biotinylated, BA-1000, RRID:AB_2313606) was used at a dilution of 1:200. All sections were processed in one batch. Images were captured using the 5x and 63x objectives with an inverted Zeiss microscope under the same settings for all slices.

### Double fluorescent-Immunohistochemistry

Double-immunostainings were performed following the previously described protocol [28], with the following modifications. We utilized 70µm thick vibratome slices for the experiment, and all conditions were conducted in the same batch. Sections were blocked with 1x ROTI®Immuno Block for 1 hour at room temperature (RT). The antibody dilutions used were as follows: anti-FoxP1, anti-FoxP2, and anti-PV at a dilution of 1:1000, and anti-SETBP1 at a dilution of 1:500. For the secondary antibodies, we used donkey anti-goat (Invitrogen, Alexa 488, A11055, RRID:AB_2534102), donkey anti-mouse (Invitrogen, Alexa 488, A21202, RRID:AB_141607), and donkey anti-rabbit (Invitrogen, Alexa 488, A21202, RRID:AB_141607), all at a dilution of 1:200. To visualize nuclei, all sections were counterstained with 4′,6-Diamidin-2-phenylindol (DAPI, Serva). Co-localization data were analyzed by manually counting co-localization in 200x200µm confocal images using the cell counter tool of the Fiji software package [44]. The co-localization of zebra finch SETBP1 with either FoxP1, FoxP2, or PV was analyzed in a total of NS (n=5), US (n=7), DS (n=6), and juvenile (n=5) samples.

### Confocal imaging and quantification of intensities from fluorescent-Immunohistochemistry

Z-stacks of zebra finch SETBP1 and co-localization experiments in Area X were acquired with a SP8-1 confocal microscope (Leica). All microscope settings were kept constant for all conditions and slices. Scans of all conditions were performed using a 40x lens with an image size of 1024×1024 pixels and a z-stack size of 1μm. The acquired images were processed using the Fiji software package [44]. For each condition, we quantified an area of 200x200µm randomly placed in the acquired image. The Rolling Ball Background Subtraction plugin was utilized to subtract background, and only nuclei with a mean gray value (MGV) >25 were quantified. We measured the mean gray values of nuclear SETBP1, or nuclear SETBP1 and nuclear FoxP2 in SETBP1+/FoxP2+ confirmed nuclei, by positioning a circle of 6μm in diameter in the center of the positive nucleus. The intensity of the SETBP1-dependent fluorescence was analyzed in a total of 637 neurons for NS (n=5 birds), 994 neurons for US (n=7 birds), 909 neurons for DS (n=6 birds), and 660 neurons for juveniles (n=5 birds). For SETBP1+/FoxP2+ co-localization, the fluorescence intensity of zebra finch SETBP1 and FOXP2 was analyzed in each neuron. We analyzed 60 neurons for NS (n=3 birds), 80 neurons for US (n=4 birds), 80 neurons for DS (n=4 birds), and 60 neurons for juveniles (n=3 birds).

### Cloning of FoxP2 Promoters

Genomic DNA from the blood of an adult male zebra finch was used as the template to amplify the promoter regions upstream of FoxP2 TSS1 and TSS2. We downloaded 380 kB of the PacBio [45] sequence upstream of the first coding exon of FoxP2 and aligned the sequences upstream of TSS1 and TSS2 from humans to it [29]. Primers were designed to amplify the entire genomic region upstream of TSS1 of the zebra finch, spanning 2,181 bp (5’-gatGCTAGCGGCATTTCACTCAGCCTCAT-3’ forward and 5’-atcAGGCCTCCCGGGTACTTTTTCCAGA-3’ reverse), and TSS2, spanning 2,501 bp (5’-gatGCTAGCTGGGTAAAATGAGAATGTAGGC-3’ forward and 5’-atcAGGCCTTCCCAGACTGATGGCATTTT-3’ reverse). Both forward primers have an NheI restriction site, and the reverse primers have a StuI restriction site. Platinum SuperFi II DNA polymerase (Invitrogen 12361010) was used to amplify the fragments. The resulting polymerase chain reaction (PCR) product was examined on an agarose gel, cleaned from nucleotides with the Qiaquick PCR purification kit (Qiagen, Chatsworth, CA), and cloned into the pGL4.13 vector using the mentioned restriction sites. We sequenced at least four clones for each of the TSS. A consensus sequence was deposited in NCBI, and the accession numbers of both TSS are OR270935 and OR270936.

### Luciferase Promoter Reporter Assays for FoxP2 promoters

Luciferase assays were conducted in HEK293 cells following the previously described protocol [42, 46]. Five luciferase assays for each TSS were performed, with each assay conducted after an independent transfection. Within each assay, triplicates were run, meaning three wells contained the same transfection reagents and quantity of cells. The mean of the triplicates was utilized for statistical analysis. Each plate was measured once in the ELISA reader. Luminescence was measured using the Dual Glo Luciferase Kit (Promega) following the manufacturer’s protocol in an ELISA plate reader (Tecan, GENios; Switzerland). The mean background from untransfected wells was subtracted from all other wells. Luciferase results are presented as the mean normalized Luciferase activity relative to the control from five independent assays.

### Statistics

GraphPad and Python were utilized to generate all graphs and analyze data. A significance level of p < 0.05 was set for all tests. For the analysis of co-localization, GraphPad was used. For the analysis of SETBP1 intensities, the mean intensity of each bird was analyzed using ANOVA followed by Tukey’s multiple comparison test. Density plots were generated using the kernel density estimation method in Python. For the analysis of correlation between singing and SETBP1 intensities, as well as the correlation between zebra finch SETBP1 and FoxP2 intensities, Spearman correlation analysis was employed. For the Luciferase assays, a one-way ANOVA followed by Tukey’s multiple comparison test was conducted.

## Data availability statement

The sequences of the four zebra finch SETBP1 isoforms were deposited to NCBI (NCBI accession numbers: OR257526-OR257529).

## Author contributions

EM and MMKW conceived general idea and together with DG, SLPdeC and ND designed all experiments. EM was involved in SETBP1 cloning. EM and SLPdeC searched for antibodies. DG, SLPdeC and EM generated all zebra finch conditions and co-localization and intensity data. EM scanned all confocal images. ND generated all DAB staining and took all DAB images. EM and MMKW were involved in cloning of TSSs of FoxP2 and luciferase assays. EM, DG and MMKW generated all figures. EM, PN and DG analyze the data and conducted statistical analysis. EM took lead in writing the manuscript. All authors provided critical feedback and helped shape the manuscript.

## Acknowledgments

Ezequiel Mendoza was supported FU Starting Grant, Nr. 43 mit FK-Beschluss vom 06.02.2023. Maggie MK Wong is supported by Max Planck Society. We sincerely thank Ursula Kobalz and Nshdejan Arpik for their invaluable technical assistance. We would like to express our gratitude to Prof. Constance Scharff, Prof. Katja Nowick, and Sophie Holtz for their invaluable assistance in proofreading and providing critical feedback on the manuscript. Additionally, we would like to acknowledge the support of the Core Facility BioSupraMol, which is funded by the DFG.

## Additional Information (Competing Interests Statement)

The authors declare no competing interests.

