## Supplemental material for "Expression and regulation of SETBP1 in the song system of male zebra finches (*Taeniopygia guttata*) during singing"

Abbreviated title: SETBP1 in male zebra finches

Author names and affiliations, including postal codes
Grönberg, Dana[1]https://orcid.org/0009-0004-3231-778X; Pinto de Carvalho, Sara Luisa[1]https://orcid.org/0009-0005-8457-1760; Dernerova, Nikola[1]https://orcid.org/0009-0006-3589-5782; Norton, Phillip https://orcid.org/0000-0002-3137-2582[2]; Wong, Maggie M. K. https://orcid.org/0000-0002-9438-0141[3, #] and *Mendoza, Ezequiel https://orcid.org/0000-0003-4963-519X [1, #].
[1] Institut für Verhaltensbiologie, Freie Universität Berlin, 14195 Berlin, Germany
[2]Humboldt-Universität zu Berlin, Institute for Theoretical Biology, Philippstr. 13, Haus 4 (Ostertaghaus), 10115 Berlin, Germany
[3] Language and Genetics Department, Max Planck Institute for Psycholinguistics, 6500AH Nijmegen, the Netherlands
### Contribute equally

**Supplementary table 1**. Amino acid comparison of zebra finch and human SETBP1 protein domains

| Domain | aa (IsoA) | aa human protein (Morgan et al 2021) | % similarity |
|---|---|---|---|
| AT-Hook domain 1 | 625-637 | 584–596 | 100 |
| AT-Hook domain 2 | 1058-1070 | 1016–1028 | 100 |
| AT-Hook domain 3 | 1490-1502 | 1451–1463 | 85 |
| HCF1 binding motiv | 1033-1036 | 991–994 | 100 |
| NLS 1 | 505-520 | 462–477 | 56 |
| NLS 2 | 1409-1423 | 1370–1384 | 53 |
| NLS 3 | 1422-1438 | 1383– 1399 | 65 |
| SET-binding domain | 1332-1527 | 1292–1488 | 80 |
| SKI homologous region | 748-959 | 706– 917 | 92 |
| PEST 1 | 1-13 | 1-13 | 92 |
| PEST 2 | 307-318 | 269–280 | 50 |
| PEST 3 | 589-601 | 548– 561 | 69 |
| PEST 4 | 719-731 | 678–689 | 92 |
| PEST 5 | 848-872 | 806–830 | 96 |
| PEST 6 | 1541-1565 | 1502–1526 | 92 |
| PPLPPPPP 1 | 1563-1570 | 1520-1527 | 100 |
| PPLPPPPP 2 | not present | 1528-1535 | / |
| PPLPPPPP 3 | not present | 1536-1543 | / |

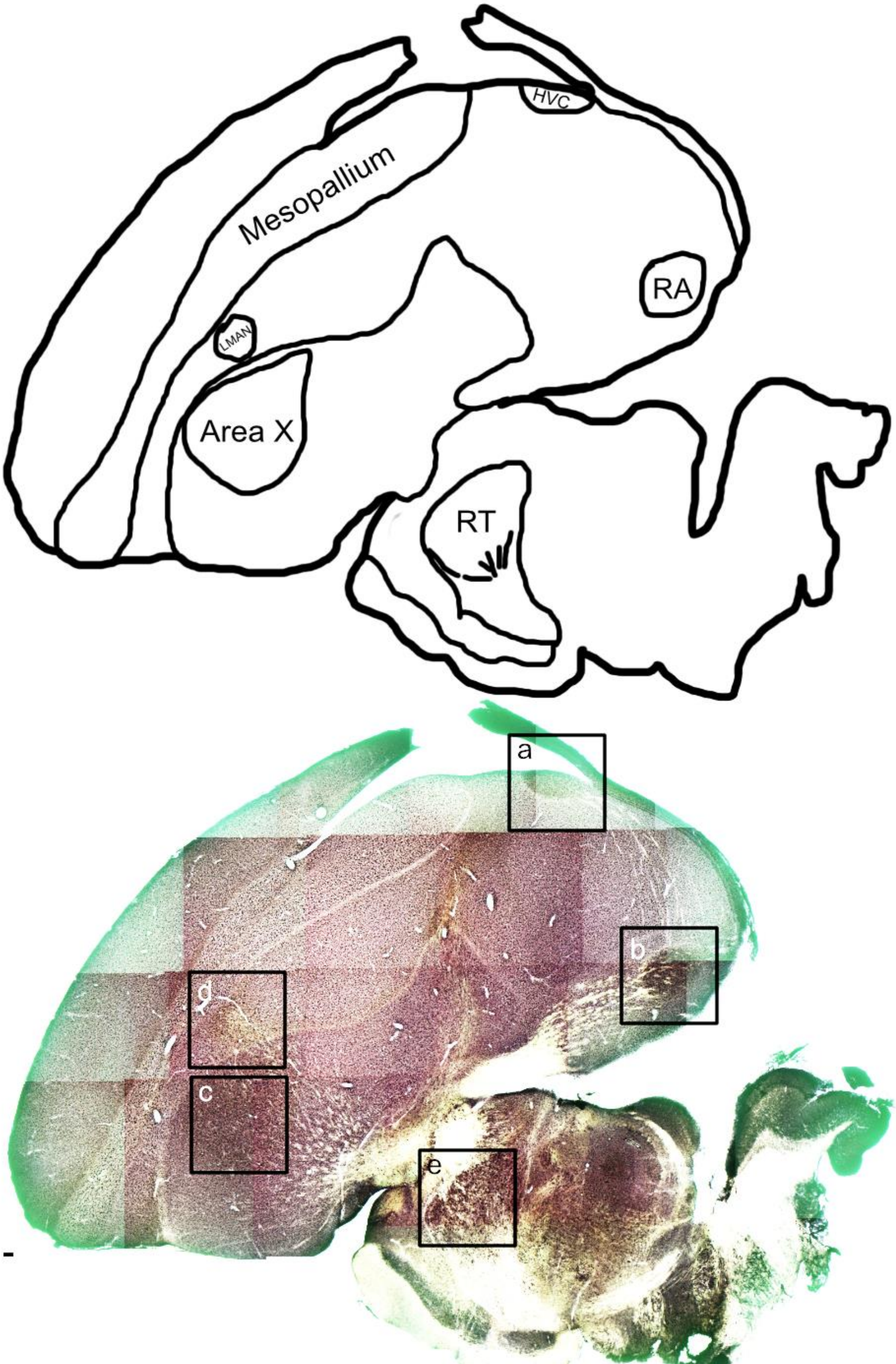

Mesopallium
HVC
RA
LMAN
Area X
RT
a
b
c
d
e

**Supplementary figure 1.-** Expression pattern of zebra finch SETBP1 protein after DAB immunodetection in a non-singing adult male (related to Figure 4). Upper part shows a schematic representation of the brain slice pointing out the regions where higher magnification photos were taken or are mentioned in the article. Names and locations of song system nuclei are shown: HVC (4a), nucleus robustus arcopallii (RA, 4b), Area X (4c), nucleus lateralis magnocellularis nidopallii anterioris (LMAN, 4d), and nucleus rotundus (RT, 4e), as well a mesopallium the region where figure 3 photos were taken. Lower part is a representative brightfield photomicrographs of a sagittal section detected with an antibody against SETBP1 and revealed with nickel enhanced DAB staining. HVC, RA, and Area X show a darker staining than the surroundings, LMAN has a weaker staining than surrounding. Higher magnification photos of figure 4 were taken from this slice (squares show the regions where the higher magnification photos were taken) and an adjacent slice was the NPC also shown in Figure 4. Scale bar = 100µm.

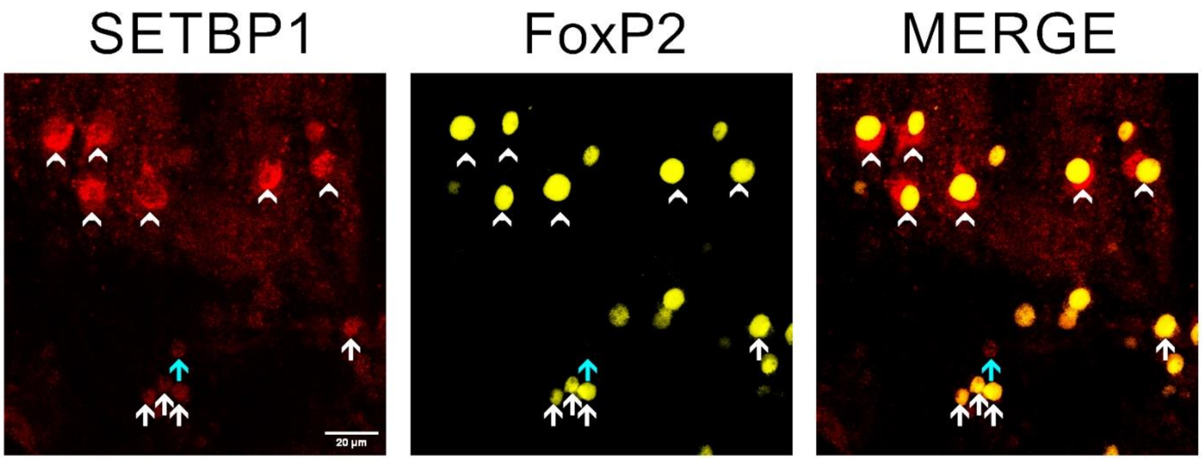


**Supplementary figure 2.-** Fluorescent immunodetection of zebra finch SETBP1 showing nuclear and cytoplasmic expression in the region of nucleus rotundus (RT) and co-localization with FoxP2. Zebra finch SETBP1 (color-coded in red) and FoxP2 (pseudo-color-coded in yellow) immunostainings showing co-localization examples in RT. Zebra finch SETBP1 is mostly expressed cytoplasmic in RT region (arrowheads) and co-localize with FoxP2, in contrast to the surrounding tissue where zebra finch SETBP1 is nuclear and may be co-localized with FoxP2 (white arrows) or not (blue arrows). Scale bar = 20µm.

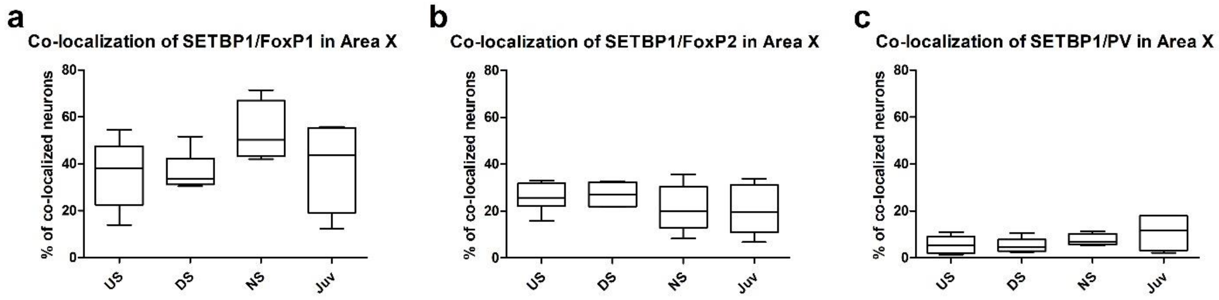


**Supplementary figure 3.-** Quantification of the Co-localization of zebra finch SETBP1 with FoxP1, or FoxP2 or parvalbumin in Area X in different conditions. a) Box plots showing the mean percentage of zebra finch SETBP1 and FoxP1 co-localizing neurons in Area X under different conditions (US, DS, NS and juvenile). b) Box plots showing the mean percentage of zebra finch SETBP1 and FoxP2 co-localizing neurons in Area X in different conditions (US, DS, NS and juvenile). c) Box plots showing the mean percentage of zebra finch SETBP1 and PV co-localizing neurons in Area X in different conditions (US, DS, NS and juvenile). Sample size (number of birds): mean of US=7, DS=6, NS=5 and Juveniles=5. No statistically significant differences between the experimental conditions were detected, one-way ANOVA, Tukey's Multiple Comparison Test.
